# Disrupted structural connectivity in ArcAβ mouse model of Aβ amyloidosis

**DOI:** 10.1101/2020.04.27.064634

**Authors:** Md. Mamun Al-Amin, Joanes Grandjean, Jan Klohs, Jungsu Kim

**Affiliations:** Department of medical and molecular genetics, Indiana University, USA; Stark Neurosciences Research Institute, Indiana University School of Medicine, Indiana University, USA; Department of Radiology and Nuclear Medicine, Donders Institute for Brain, Cognition, and Behavior, Donders Institute, Radboud University Medical Center, Nijmegen, Netherlands; Institute for Biomedical Engineering, University of Zurich & ETH Zurich, 8093 Zurich, Switzerland

**Keywords:** Alzheimer’s Disease, white matter, neurofilament, connectivity, perineuronal net, Aβ

## Abstract

Although amyloid beta (Aβ) deposition is one of the major causes of white matter (WM) alterations in Alzheimer’s disease (AD), little is known about the underlying basis of WM damage and its association with global structural connectivity and network topology. We aimed to dissect the contributions of WM microstructure to structural connectivity and network properties in the ArcAβ mice model of Aβ amyloidosis.

We acquired diffusion-weighted images (DWI) of wild type (WT) and ArcAβ transgenic (TG) mice using a 9.4 T MRI scanner. Fixel-based analysis (FBA) was performed to measure fiber tract-specific properties. We also performed three complementary experiments; to identify the global differences in structural connectivity, to compute network properties and to measure cellular basis of white matter alterations.

Transgenic mice displayed disrupted structural connectivity centered to the entorhinal cortex (EC) and a lower fiber density and fiber bundle cross-section. In addition, there was a reduced network efficiency and degree centrality in weighted structural connectivity in the transgenic mice. To further examine the underlying neuronal basis of connectivity and network deficits, we performed histology experiments. We found no alteration in myelination and an increased level of neurofilament light (NFL) in the brain regions with disrupted connectivity in the TG mice. Furthermore, TG mice had a reduced number of perineuronal nets (PNN) in the EC.

The observed FDC reductions may indicate a decrease in axonal diameter or axon count which would explain the basis of connectivity deficits and reduced network efficiency in TG mice. The increase in NFL suggests a breakdown of axonal integrity, which would reduce WM fiber health. Considering the pivotal role of the EC in AD, Aβ deposition may primarily increase NFL release, damaging PNN in the entorhinal pathway, resulting in disrupted structural connectivity.

## Introduction

Alzheimer’s disease (AD) is a progressive cognitive disorder, affecting 5.6 millions of people in the United States with an estimated caregiver cost of nearly $244 billion (Alzheimer’s Association, 2020). AD negatively impacts on patient’s daily life by deteriorating cognitive components; attention, memory, reasoning, thoughts and decision making. AD pathology has been reported as a series of progressive events that begins with the aggregation and deposition of Amyloid-β (Aβ, a metabolite of amyloid precursor protein (APP)) followed by accumulation of hyper-phosphorylated tau proteins, impaired neuronal function and neuronal death (Jack et al., 2013). Cerebral amyloid and tau burden is strongly associated with the atrophy of gray matters (GM) (Sepulcre et al., 2016) and hyperintensity of white matters (WM) (Weaver et al., 2019).

Over the past decade, multimodal neuroimaging techniques have been used to unravel the impact of Tau and Aβ accumulation on both GM and WM pathology. Functional MRI (fMRI) studies show reduced activity of functional networks and connectivity (Sepulcre et al., 2017, Pereira et al., 2019, Son et al., 2017), while diffusion-weighted MRI (DWI) studies demonstrate impaired WM integrity in AD (Agosta et al., 2011, Acosta-Cabronero et al., 2010). To determine the WM properties, Tract-based spatial statistical (TBSS) analytical technique has been used over the years in various disease contexts. However, TBSS only measure the widespread alterations of WM properties and does not address fiber-specific properties, including fiber density (FD) or fiber cross-section (FC). Newer approaches, such as neurite orientation dispersion and density imaging (NODDI), have been used to profile neurite density indexes (NDI) and orientation dispersion indexes (ODI) (Wang et al., 2019) in the mouse models of Aβ amyloidosis (Colon-Perez et al., 2019) and tau pathology (Colgan et al., 2016). However, a WM bundle comprises of many axonal fibers and aforementioned techniques do not allow study of fiber-specific WM properties; such as FD, FC and fiber density plus cross-section (FDC). Nevertheless, reduction in FD, FC and FDC are seen in patients with AD and mild cognitive impairment (Mito et al., 2018). To date, there is a lack of knowledge on whether fiber-specific WM properties are compromised in AD.

WM facilitates communications between brain regions through complex neuronal networks. Network-based theories suggest that such structural networks are selectively vulnerable in AD (Andreotti et al., 2017). Disrupted functional and structural connectivity has been extensively observed in animal models of Aβ amyloidosis. These includes an altered resting-state functional network and compromised WM indices have shown in TgF344 rats (Anckaerts et al., 2019), 5XFAD mice (Kesler et al., 2018), APP/PS1 mice (Shah et al., 2013, Bero et al., 2012), ArcAβ mice (Grandjean et al., 2014, Grandjean et al., 2016), aging apoE4 and apoE-KO mice (Zerbi et al., 2014) and PSAPP mice (Grandjean et al., 2016), although not in E22A*Δ*β mice (Grandjean et al., 2016). However, these studies did not measure the fiber-specific tract properties mentioned earlier. In addition, it is less clear whether structural connectivity is altered in the mouse model of Aβ amyloidosis. As WM lesions contribute to the progression of AD (Shao et al., 2012) and lead to impaired structural connectivity (Vogel et al., 2019) in humans, we aimed to assess structural connectivity in the ArcAβ mice. Previous study showed an increased white matter property, such as axial diffusivity in minor forceps and external capsule, in the ArcAβ mice (Grandjean et al., 2016). Therefore, we hypothesized an altered properties of white matter fiber tract in the ArcAβ transgenic mice.

The underlying basis of WM lesions following Aβ accumulation has been studied in rodents. WMs consisting of nerve fibers (bundles or axons) enriched with neurofilaments (Trojanowski et al., 1986). Axonal swelling, degeneration (Wirths et al., 2007) and transport deficits (Smith et al., 2007) are shown in mouse models Aβ amyloidosis (Aβ APP/PS1ki). Besides myelin, the level of neurofilament in cerebrospinal fluid (CSF) and plasma associates with axonal injury (Menke et al., 2015). Neurofilament is expressed in the WM tracts (Bergman et al., 2016) and categorized into; light, medium and heavy. Neurofilament light (NFL) is the most abundant in the brain and is required for the radical growth of axons during the development (Eyer and Peterson, 1994). Neurofilaments accumulate specifically in the cell bodies and proximal areas of neurons during neurological disorders (Liu et al., 2009). The concentrations of NFL is increased both in the plasma (Lin et al., 2018) and CSF (Jin et al., 2019, Zetterberg et al., 2016) in AD. The level of CSF NFL increases as a function of age (Bacioglu et al., 2016) in mice model of Aβ amyloidosis, while NFL deficiency aggravates amyloid pathology in NFL knock out (KO) mice (Fernandez-Martos et al., 2015). Since NFL is a potential biomarker of WM damage (Sjogren et al., 2001), we hypothesized that the levels of NFL would increase in mouse model of Aβ amyloidosis.

Besides the role of myelin and NFL, the brain’s extracellular matrix (ECM) plays a potential role in maintaining the connectivity in brain networks (Bikbaev et al., 2015). Amyloid deposition damages the ECM scaffold (Ajmo et al., 2010, Sykova et al., 2005). The most well-studied type of ECM is Perineuronal Nets (PNN). This lattice-like structure envelop the soma, stabilizing interneuron activity, regulating synaptic homeostasis (Bosiacki et al., 2019) and protecting interneurons from oxidative damage (Morawski et al., 2004). The role of PNN in mouse model of Aβ amyloidosis have been inconsistent and contradictory. For instance, one study showed an unaltered PNN expression in Tg2576 mice (Morawski et al., 2010b), while others suggest that the reduction of the ECM ameliorates memory impairment in APP/PS1 mice (Vegh et al., 2014) and the degradation of PNN restores memory in TgP301s mice (Yang et al., 2015). By contrast, PNN protects neurons from Tau-associated pathological changes (Morawski et al., 2010b) and Aβ-associated neurotoxicity (Miyata et al., 2007). In summary, the role of PNN in AD pathogenesis is inconclusive.

We first aimed to analyze fiber-specific WM properties in a mouse model of Aβ amyloidosis, ArcAβ. ArcAβ mice express the human APP gene, with Swedish (K670 N & M671L) and arctic (E693G) mutations (Knobloch et al., 2007b). ArcAβ mice show a reduced synaptic function in between 3.5 to 7.5-months of age (Knobloch et al., 2007a). Aβ plaque deposition and impaired cognitive function begins at 6 months (Knobloch et al., 2007b), with vascular Aβ appearing between 9 and 15 months (Merlini et al., 2011). We began by performing whole-brain tractography to analyze differences in structural connectivity in between wild type (WT) and ArcAβ transgenic (TG) mice to identify brain regions with abnormal edges/connections. We then implemented graph theory analysis to measure the network properties. Finally, we performed histology to measure myelination, NFL, and PNN.

## Method

### Animals

Hybrid C57BL/6 and DBA/2 background mice were used to generate ArcAβ mice in the animal facilities at the Institute for Biomedical Engineering, University and ETH Zürich, Zürich, Switzerland. The animals were housed in a facility with a 12-h light-dark cycle, and a room temperature of 75°F with ad-libitum access to food and water. A total of 32 mice were included in this study. Diffusion MRI data was acquired using WT (*n* = 12) and ArcAβ TG (*n =* 8) mice (381-431 days old) [see more details in (Grandjean et al., 2016). An additional 12 mice (*n* = 6/group) that were 157-214 days old used in the immunohistochemistry. The experimental protocols were approved by the Zürich Cantonal veterinary office in Switzerland.

The mice were anesthetized (3.5% isoflurane, medetomidine and pancuronium bromide) before acquisition of *in-vivo* diffusion MRI data. Artificial ventilation (80 breaths/min; O_2_ to air = 1:4, 1.8 ml/h flow (CWE, Ardmore, USA) was administered to ensure endotracheal intubation. The level of isoflurane was gradually reduced (2%, 1.5%, 0.5%), while the levels of medetomidine was increased from 0.05 mg/kg to 0.1 mg/kg and pancuronium bromide from 0.2 mg/kg to 0.4 mg/kg. We monitored the body temperature (36.5°C ± 0.5) using a rectal thermometer probe.

### Diffusion-weighted imaging

We acquired diffusion weighted MRI data on a Bruker 94/30 Biospec spectrometer (Bruker BioSpin MRI, Ettlingen, Germany) equipped with cryogenic surface receiver coil, a resonator coil and BGA-S gradient system. The parameters of DWI data acquisition were as follows: voxel size = 0.15 × 0.13 × 0.45 mm, TR (Repetition time) = 2000 ms, TE (Echo time) = 22 ms, MS = 128 ×128, PR = 0.15 × 0.13 mm^2^, Field of view = 20 × 17.5 mm^2^. We acquired 5 volumes of *b* = 0 s/mm^2^ and *b* = 1000 s/mm^2^ with 36 directions.

### DWI data pre-processing

Briefly raw DWI image was corrected with motion correction, image inhomogeneity correction and eddy current using advanced normalization tools (Avants et al., 2009) and FSL (http://fsl.fmrib.ox.ac.uk) (Fig. 1b) (Smith, 2002). We then estimated tissue response (Fig. 1c), created a DWI mask (Chou et al., 2011) before calculating fixel orientation distribution (FOD) (Fig. 1e) in MRTRix3 (Tournier et al., 2019). Probabilistic tractography was conducted using constraint spherical deconvolution (CSD) in MRTRix3 (Tournier et al., 2007). We first registered *b* = 0 image to a previously developed atlas [combined from AMBMC (Australian Mouse Brain Mapping Consortium) and John Hopkins C57BL/6J mouse brain atlas] (Fig. 1h) (Liu et al., 2016). The registration was further cross-checked and fixed in ITK-SNAP (Yushkevich et al., 2006). The structural connectome (Fig. 1i) was constructed from 76 brain region of interest (ROI) based on our previous pipeline developed in (Al-Amin et al., 2019).

**Fig. 1.**
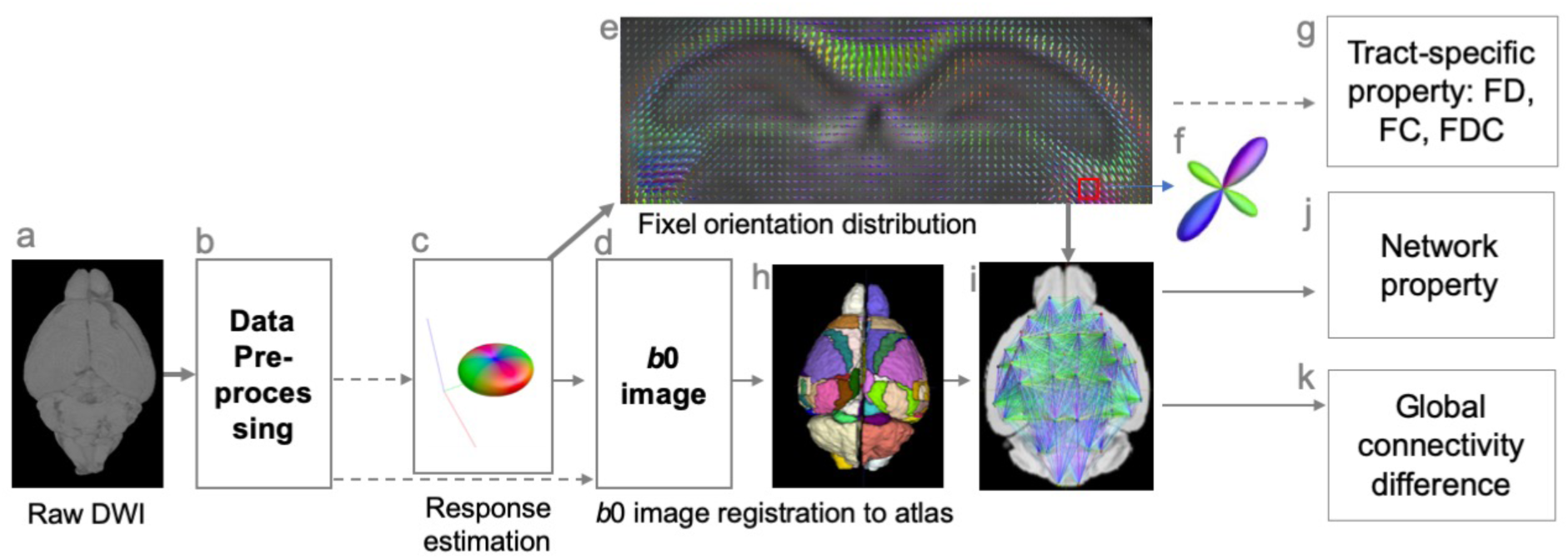
Processing pipeline of image analysis. Raw DWI data (a) was preprocessed prior to estimate tissue response (c) and fixel orientation distribution (e). The *b* = 0 image (d) was extracted from the preprocessed DWI and registered to a previously developed atlas (h) (Liu et al., 2016) to perform whole-brain probabilistic tractography. Fiber tract specific analysis was performed to measure fiber density (FD), fiber cross-section (FC) and fiber density plus cross section (FDC) (g). The structural connectomes (i) were extracted from the tractography data using 76 × 76 brain regions of interests (h). The network efficiency was measured using Gretna toolbox (Wang et al., 2015) (j). Whole-brain network difference analysis was performed with network-based statistics (k).

### Fixel-based analysis

To perform fixel-based analysis (FBA) (Fig. 1g) we followed a previous protocol (Raffelt et al., 2017) available in MRTRix3. We created a population-based FOD template based on *n* = 20 samples. To visualize all *p*-values less than 0.05, we thresholded the fixels using “mrview” (MRTRix3) at 0.95 with FWE (Family-wise error) corrected value (Fig. 1g).

### Whole-brain connectivity analysis

To identify the global differences in connectivity in TG mice, we performed network-based statistics (NBS) (Fig. 1k) (Zalesky et al., 2010). NBS is useful in controlling the FWE while identifying differences between two experimental groups. NBS used two measure: “intensity” and “extent”. In NBS “intensity” analysis indicates total test statistics value of all connections, while “extent” analysis encompasses total number of connections of network. We performed 5000 permutations from a threshold range between 2.5 to 3.5 (Al-Amin et al., 2019, Al-Amin, 2019) using two different contrasts WT > TG and TG > WT.

### Network properties analysis

The network sparsity was calculated based on 76 × 76 matrix (from 76 ROI). Since we used probabilistic tractography, there is a possibility that false connections can be produced. Therefore, we used a lower network sparsity (10% to 30%) corresponding to our threshold calculation of 1 to 10 (Fig. 4). We measured a range of graph theory parameters such as global efficiency, local efficiency, clustering coefficient, degree centrality, small-world and betweenness using GRETNA toolbox (Fig. 1j) (Wang et al., 2015).

### Immunohistochemistry

We performed transcardial perfusion with 0.1 M phosphate-buffered solution (PBS) and 4% paraformaldehyde (PFA) in PBS. We then carefully peeled the skull off the brain. The brain was fixed in 4% PFA at 4 °C for 24 h. Brains were then carefully dissected and preserved at 4 °C in 0.1 M PBS solution containing 0.02% sodium azide followed by paraffin embedding. We collected 10 µm thick coronal slices using a rotary microtome. A one-in-ten series of the coronal sections were stained to label NFL, myelin basic protein (MBP) and PNN-expressing neurons.

To retrieve antigens, we immersed the brain sections in Antigen Recovery Solution (citrate buffer solution containing of 10 mM Sodium Citrate, 0.05% Tween 20, pH 6.0) at 70°C for 30 min followed by three washes in PBS. We then used a blocking solution (2.5% normal goat serum, 0.1% Triton X-100 and 10% sodium azide in PBS) to block the sections at room temperature for 1 h. Sections were then incubated with primary antibodies; rabbit anti-NFL (1:500) (C28E10, a gift from Cell-signaling, Chicken anti-MBP (1:1000) (a gift from EnCor Biotechnology Inc) and biotinylated WFA (*Wisteria floribunda* agglutinin) (B-1355, Vector Laboratories, USA) (1:2000) at room temperature for 48 h. We used secondary antibodies that were conjugated with fluorescent markers such as Alexa fluor^®^ 488 anti-rabbit (1:1000) (Invitrogen), Alexa fluor^®^ anti-chicken (1:1000) (Invitrogen)and Alexa Fluor^®^ 488 Streptavidin (1:1000) (Jackson ImmunoResearch Laboratories Inc.) at room temperature for 24 h. The sections were then mounted with Vectashield containing DAPI (Vector laboratories, USA).

### Microscopy and image analysis

We captured fluorescence images on a Leica Slide Scanner microscope using a 20x objective. Cells that immunolabelled with NFL, MBP, WFA and DAPI were included in the imaging. The intensity of NFL, MBP and WFA staining was analyzed. In addition, the number of WFA-positive cells were counted from the dorsal hippocampal regions, entorhinal cortex (EC), secondary visual cortex (V2), secondary visual cortex mediolateral(V2ml), and white matter (WM) (Fig. 7A). Image analysis was performed in Fiji (ImageJ) (Schindelin et al., 2012). For each mouse we analyzed three coronal sections in matching anatomical locations (starting 2.30 mm - 2.80 mm posterior to bregma).

## Statistical Analysis

We conducted statistical analyses using SPSS (Version 24.0) and GraphPad PRISM (Version 8.0). We performed a two-way ANOVA followed by the Bonferroni post hoc test when the P-value was < 0.05, to measure the difference between groups. We considered the effect of APP mutation (WT vs TG) and brain region (Ent, Hip, V2, V2ml, WM) as an independent variables and intensity of NFL, MBP, PNN and PNN count as dependent variables. Further post-A *p*-value below 0.05 was considered significant. Data are presented as mean ± SEM.

## Results

### Decreased fiber tract-specific property in ArcAβ transgenic mice

Fixel-based analysis revealed a significant (FWE corrected *p*-value *≤* 0.05) reductions of streamline segments in TG mice. The streamline segments were associated with fiber density (Fig. 2a), fiber bundle cross-section (Fig. 2b) and fiber density plus fiber bundle cross-section (FDC) (Fig. 2c). The color gradient represents the magnitude of the reduction in the TG mice compared to their counterpart wild type group. Observed decreased FD, FC and FDC were shown in the right hemisphere, particularly in the corpus callosum, hippocampal commissure, retrosplenial cortex, dorsal hippocampus and thalamus. Interestingly, when FD and FC were considered together, the FDC metric had shown a larger difference in the TG mice.

**Fig. 2.**
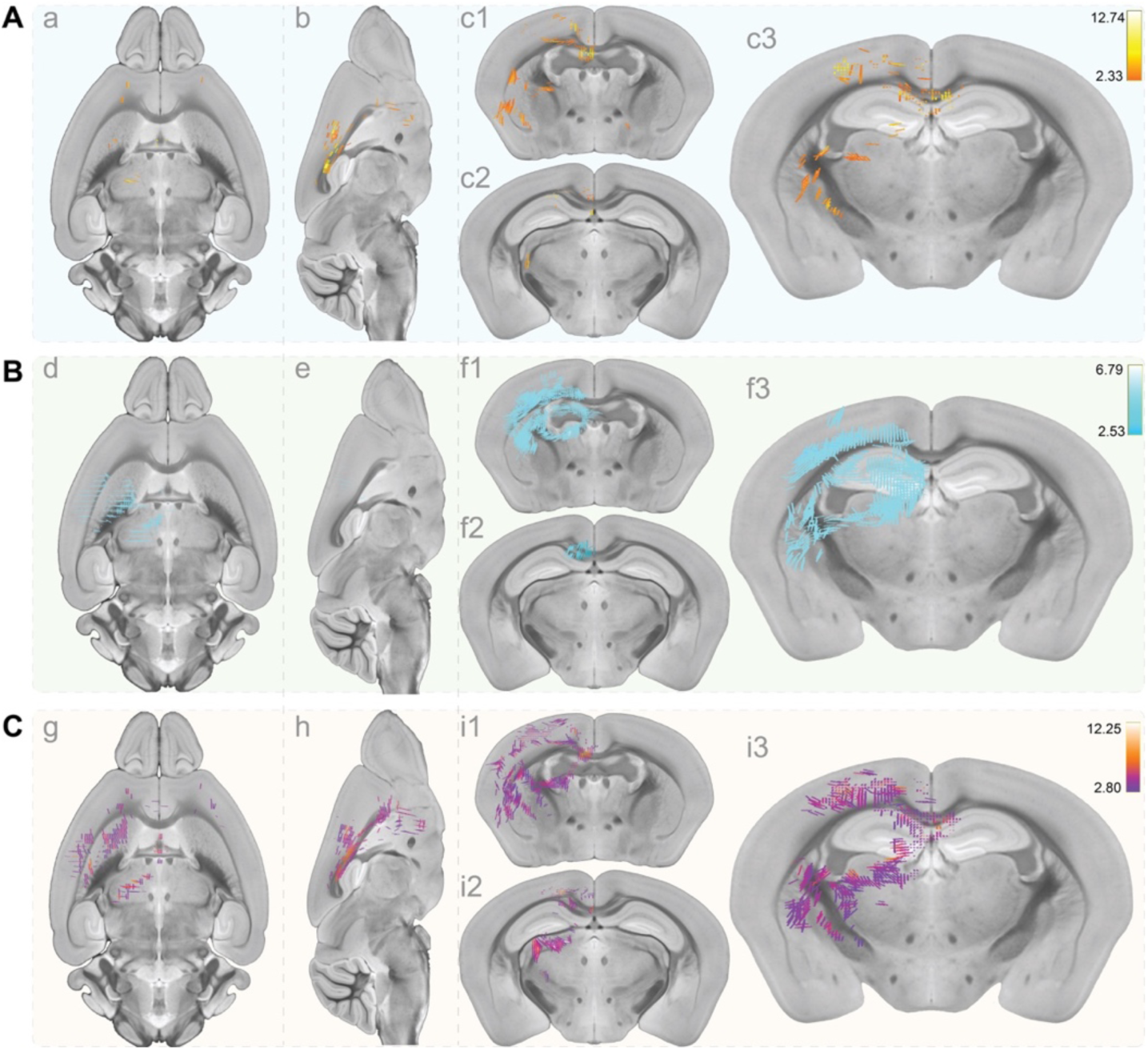
Transgenic mice exhibits a decrease in fiber-specific properties. Transgenic mice display a reduction in fiber density (FD) (row A), fiber bundle cross-section (FC) (row B), and fiber density plus fiber bundle cross-section (FDC) (row C). The streamlines were found following Family wise error (FWE) correction at *p* < 0.05 and overlaid on the AMBMC atlas (Ullmann et al., 2014). The color gradient on the right side, indicates the magnitude of the effect size. The result shows based on the contrast of WT > TG statistical analysis. No streamlines were shown with the contrast TG > WT. The groups are WT (*n* = 12), Wild Type, TG (*n* = 8), Transgenic, TG ArcAβ mice. Axial (left column), Coronal (right column) and Sagittal (middle column) views of the brain.

### Disruption in whole-brain structural connectivity in ArcAβ transgenic mice

To identify differences in whole-brain structural connectivity differences in between ArcAβ WT and TG mice, we performed network-based statistics analysis. We observed the lowest *p*-value in both network intensity (*p* = 0.029) and network extent (*p* = 0.048) at a threshold of *t* = 3.4, so this threshold was used for NBS. NBS analysis revealed brain network deficits in the TG mice compared to WT (Fig. 3). Connection deficits were observed in five brain regions (or nodes) in the right hemisphere: entorhinal cortex (EC.R), periaqueductal grey (Pag.R); subiculum (Sub.R); mediolateral cortex (V2ml.R) and secondary (V2.R) visual cortex. Based on the number of edges, entorhinal cortex (3 edges) was the biggest module in the disrupted network.

**Fig. 3.**
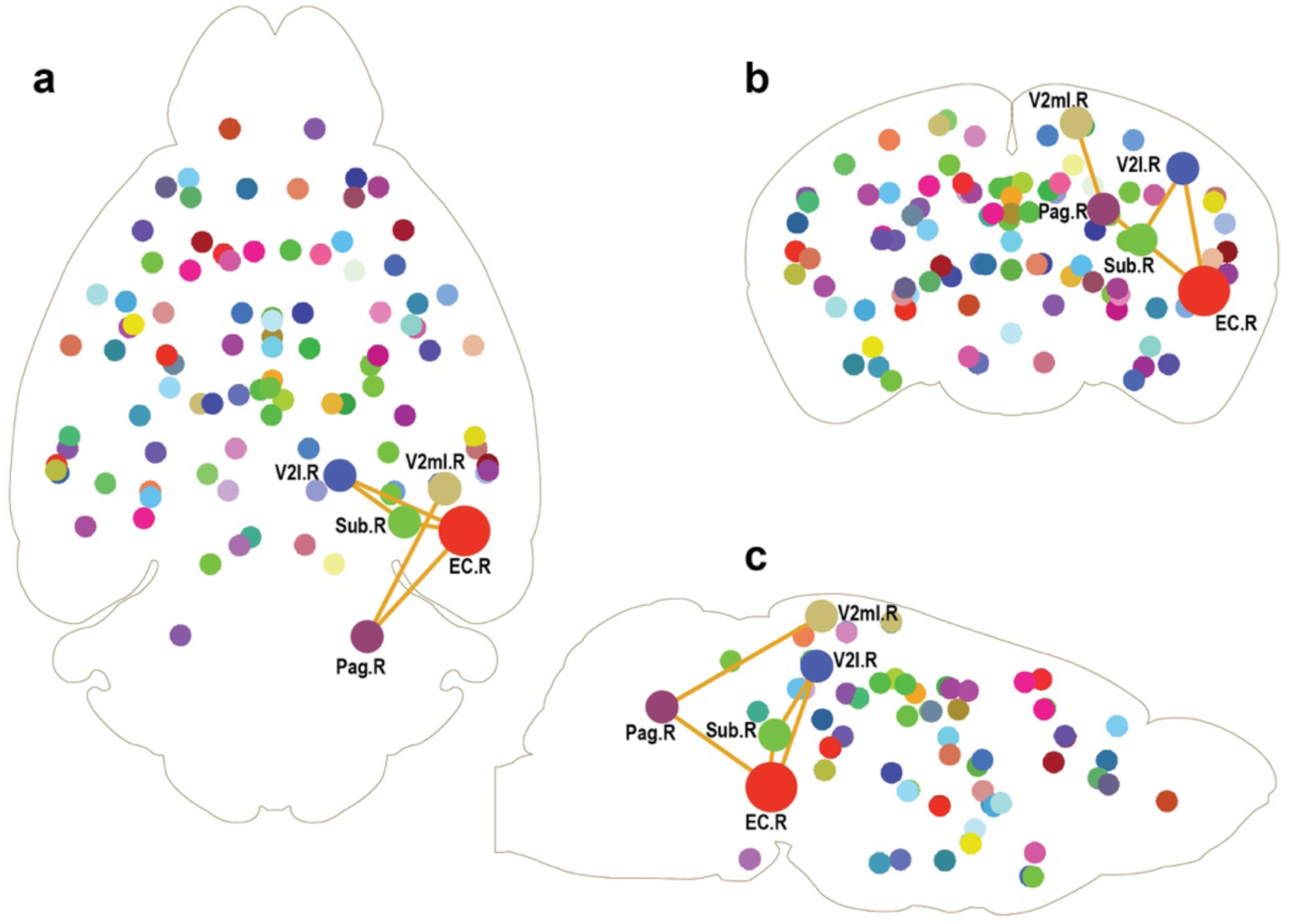
ArcAβ transgenic mice displays a disruption of structural connectivity centered to the right entorhinal cortex. At a statistical threshold of *t* = 3.4, this network shows the highest statistical significance in terms of extent and intensity. The disrupted connectivity (edges) were shown in five nodes: *Ent.R*, Entorhinal cortex (right); Pag.R, periaqueductal grey (right); Sub.R, Subiculum (right); V2ml.R, mediolateral cortex (right); V2l.R, secondary visual cortex (right). Network based statistics results are displayed in axial (a), coronal (b) and sagittal (c). The groups are WT (*n* = 12), Wild Type; TG (*n* = 8), Transgenic ArcAβ mice.

### Decreased network properties in ArcAβ transgenic mice

Since structural connectivity is the basis of brain networks, we hypothesized that the disrupted structural connectivity would alter network properties in the TG mice. Therefore, to measure network properties, we performed network-wise comparison by applying graph theory on the weighted structural network. We used different network densities (or threshold) starting from 10% to 30% (corresponding threshold 1 to 10 in Fig. 4). We also calculated the “area under the curve” (AUC) of each network to provide a scalar that are independent to any specific threshold. Independent sample *t*-tests were conducted on the “area under the curve” (AUC) of the net-work measures between the WT and ArcAβ TG mice (Fig. 4A). Transgenic mice had significant reductions in degree centrality (aDc) (*t*13 = 2.21, *p* = 0.045) (Fig. 4Ab), and local efficiency (aEloc) (*t*_13_ = 2.26, *p* = 0.041) (Fig. 4Ad), but not betweenness centrality (aBc) (*t*_13_ = 0.55, *p* = 0.938) (Fig. 4Aa), global efficiency (aEg) (*t*_13_ = 2.14, *p* = 0.127) (Fig. 4Ac), small-world clustering coefficient (aCp) (*t*_13_ = 1.77, *p* = 0.095) (Fig. 4Ae), small-world path length (aLp) (*t*_13_ = 1.77, *p* = 0.69) (Fig. 4Af), nodal clustering coefficient (aNCp) (*t*_13_ = 0.498, *p* = 0.095) (Fig. 4Ag) and nodal shortest path length (aNLp) (*t*_13_ = 1.96, *p* = 0.529) (Fig. 4Ah).

**Fig. 4.**
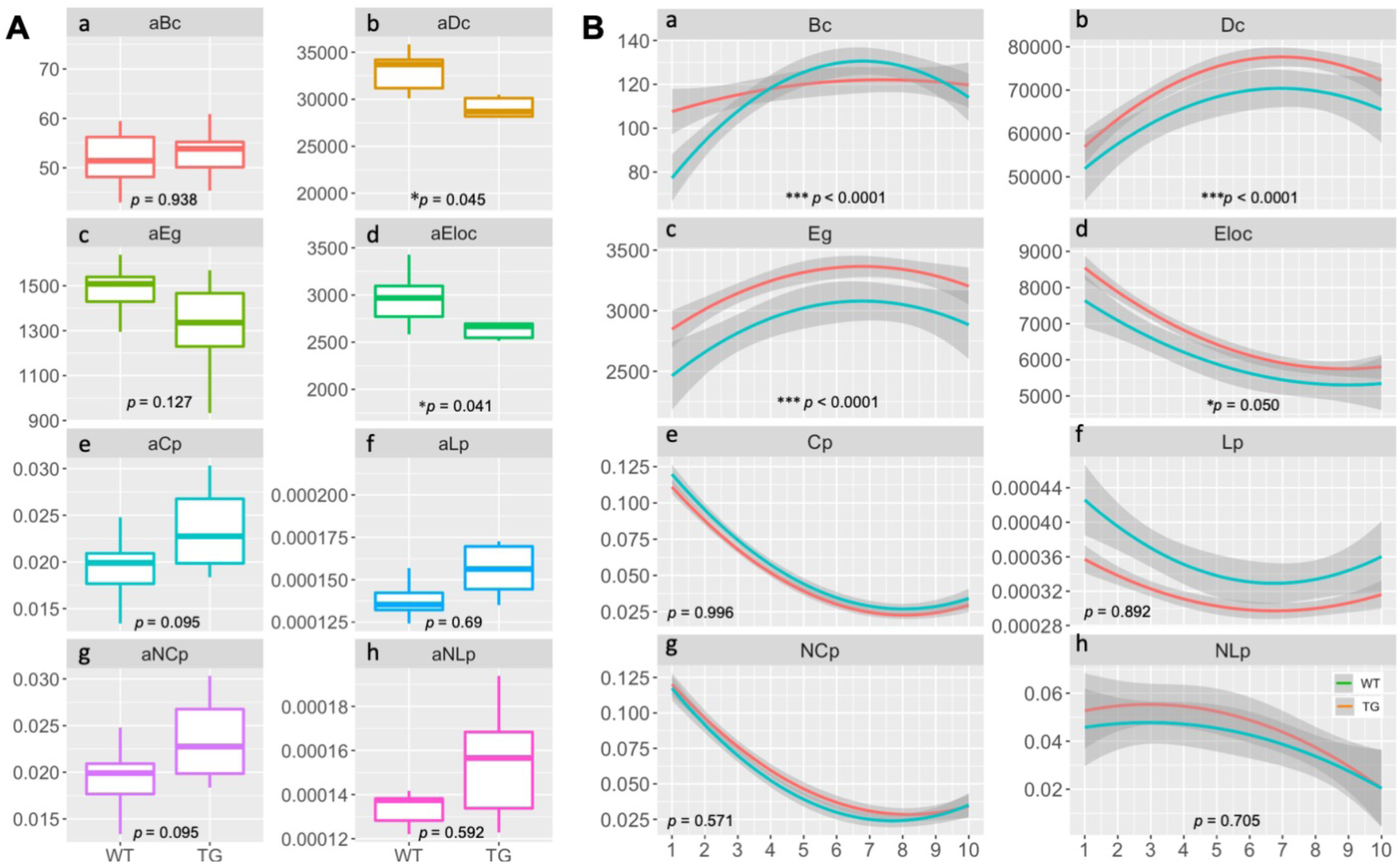
Network-wise comparison shows a reduced network topology in ArcAβ transgenic mice. Transgenic mice exhibit a decrease area under the curve (AUC) (A) of degree centrality (aDc) (Ab) and local efficiency (aEloc) (Ad) compared to the wild type mice. Transgenic mice also display decrease betweenness centrality (Ba) degree centrality (Bb), global and local efficiency (Bc and Bd) at different network densities (B). Values are means ± SEM. The groups are WT (*n* =12), Wild Type; TG (*n* = 8), Transgenic ArcAβ mice.

We also performed a two-way ANOVA (threshold x network parameters) to identify the main effect of genotype on network parameters affected by genotype at a range of different network densities. There was a significant interaction between Genotype and Threshold (*F*_9,126_ = 16.95; *p* < 0.0001) and a main effect of Genotype (*F*_1,14_ = 0.003; *p* = 0.951) on Betweenness centrality (Fig. 4Ba). There was also a significant interaction between Genotype and Threshold (*F*_9,126_ = 3.91; *p* < 0.001) and main effect of Genotype (*F*_1,14_ = 5.073; *p* < 0.001) on Degree centrality (Fig. 4Bb).

Likewise, there was a significant interaction between Genotype and Threshold (*F*_9,126_ = 5.27; *p* < 0.0001) and main effect of Genotype (*F*_1,14_ = 4.604; *p* < 0.05) on local efficiency (Fig. 4Bd). Surprisingly, there was no significant interaction between Genotype and Threshold (*F*_9,126_ = 0.900; *p* = 0.527) on global efficiency (Fig. 4Bc).

There was no significant interaction between Genotype and Threshold on small-world index Cp (clustering coefficient) (*F*_9,126_ = 0.658; *p* = 0.745) (Fig. 4Be) and Lp (path length) (*F*_9,126_ = 5.17; *p* < 0.001) (Fig. 4Bf). Likewise, there were no main effect of Genotype on nodal on clustering coefficient (*F*_1,14_ = 1.351; *p* = 0.264) (Fig. 4Bg) and nodal path length (*F*_9,126_ = 0.882; *p* = 0.543) (Fig. 4Bh).

### Increased immunoreactivity to NFL in ArcAβ transgenic mice

Having found that structural connectivity disruption is consistent with the reduced network topology, we asked whether this connectome deficit is due to alteration of fiber tracts. We used two relevant axonal markers: MBP and NFL. MBP is used to selectively measure myelination of axonal fibers, while NFL is a marker of axonal degeneration. The immunoreactivity of MBP and NFL was analyzed in the five ROIs in which connectivity was found to be impaired through network-based analysis (Fig. 3); entorhinal cortex (EC), secondary visual cortex (V2), ventromedial cortex (V2ml), hippocampus and hippocampal commissure. In addition, fixel-based analysis revealed a reduction of FD, FC and FDC in the hippocampus and hippocampal commissure (Fig. 2).

Two-way (Genotype x ROI) ANOVA analysis of NFL immunoreactivity revealed a significant interaction (*F*_4,162_ = 10.31; *p* < 0.001). In addition, there was a significant main effect of Genotype (*F* _1,162_ = 131.7; *p* < 0.001) and a significant main effect of ROI (*F*_4,162_= 8.94; *p* < 0.001) on NFL immunoreactivity. NFL immunoreactivity was significantly elevated in each brain regions in TG mice (Fig. 5e).

**Fig. 5.**
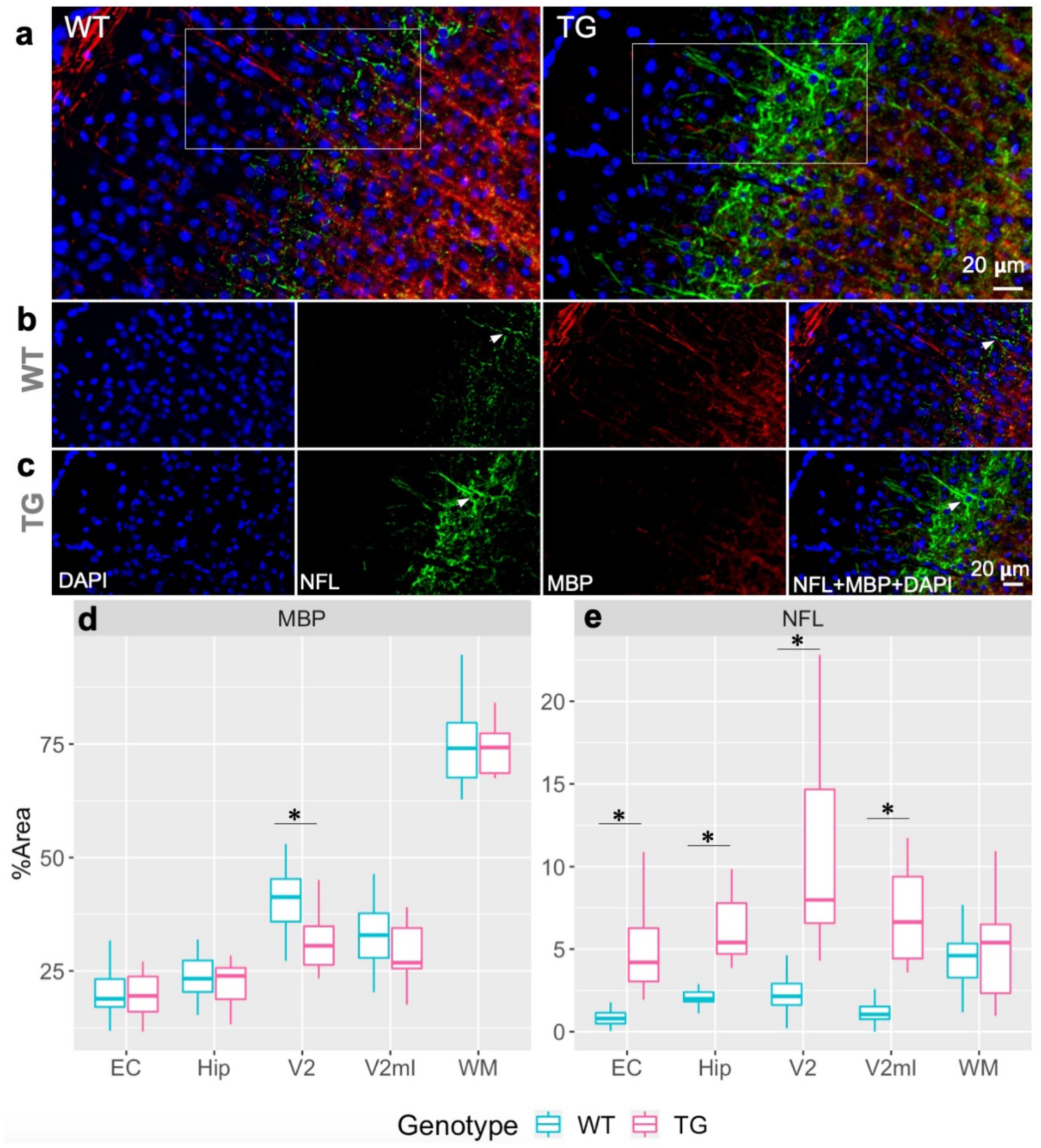
There was a significant increase in neurofilament light intensity in the entorhinal cortex (EC), hippocampus (Hip), secondary visual cortex (V2), mediolateral Secondary visual cortex (V2ml). However, reduced myelin intensity was shown only in the Secondary visual cortex (V2) (e). Triple immunostained secondary visual cortex of wildtype (WT) (a, Top Left) and transgenic (TG) (a, Top Right) mice. NFL-labelled cell (green), Myelin basic protein (MBP)-labelled dendritic processes (red) and nuclei (blue) (D). Transgenic mice higher NFL intensity (Row c) while compared to the WT mice (Row b). White arrow head indicates NFL immunostaining. Values are means ± SEM, *n* = 6/group. \*\*\**p* < 0.001 compared with the wild type (WT).

Further two-way (Genotype x ROI) ANOVA analysis on MBP intensity showed a significant main effect of Genotype (*F*_1,164_ = 9.57; *p* < 0.005) and a significant main effect of ROI (*F*_4,164_ = 304.8; *p* < 0.001). However, there was no significant interaction (*F*_4,162_ = 1.91; p = 0.111) between Genotype with ROI (Fig. 5d).

### Decrease of PNN fluorescent intensity in ArcAβ transgenic mice

To further investigate the underlying cellular basis of structural connectivity and network impairment, we immunostained neuron-specific ECM, PNN with biotinylated WFA. We calculated the number of PNN labeled cells and the intensity of WFA immunoreactivity, and performed a two-way ANOVA (Genotype × ROI). We found a significant interaction (*F*_4,165_ = 6.29; *p* < 0.001) between Genotype and ROI on PNN intensity. Moreover, there was a main effect of Genotype (*F*_1,162_ = 66.64; *p* < 0.001) and ROI (*F*_4,165_= 60.84; *p* < 0.001) on PNN intensity. The PNN immunoreactivity was significantly reduced in EC, V2 and V2ml regions in TG mice (Fig. 6d). We further analyzed WFA-labelled PNN cells and found a main effect of Genotype (*F*_1,167_ = 24.89; *p* < 0.001) and ROI (*F*_4,167_= 59.14; *p* < 0.001) but not interaction (*F*_4,167_ = 1.77; *p* < 0.136) on total number of PNN cells (Fig. 6e).

**Fig. 6.**
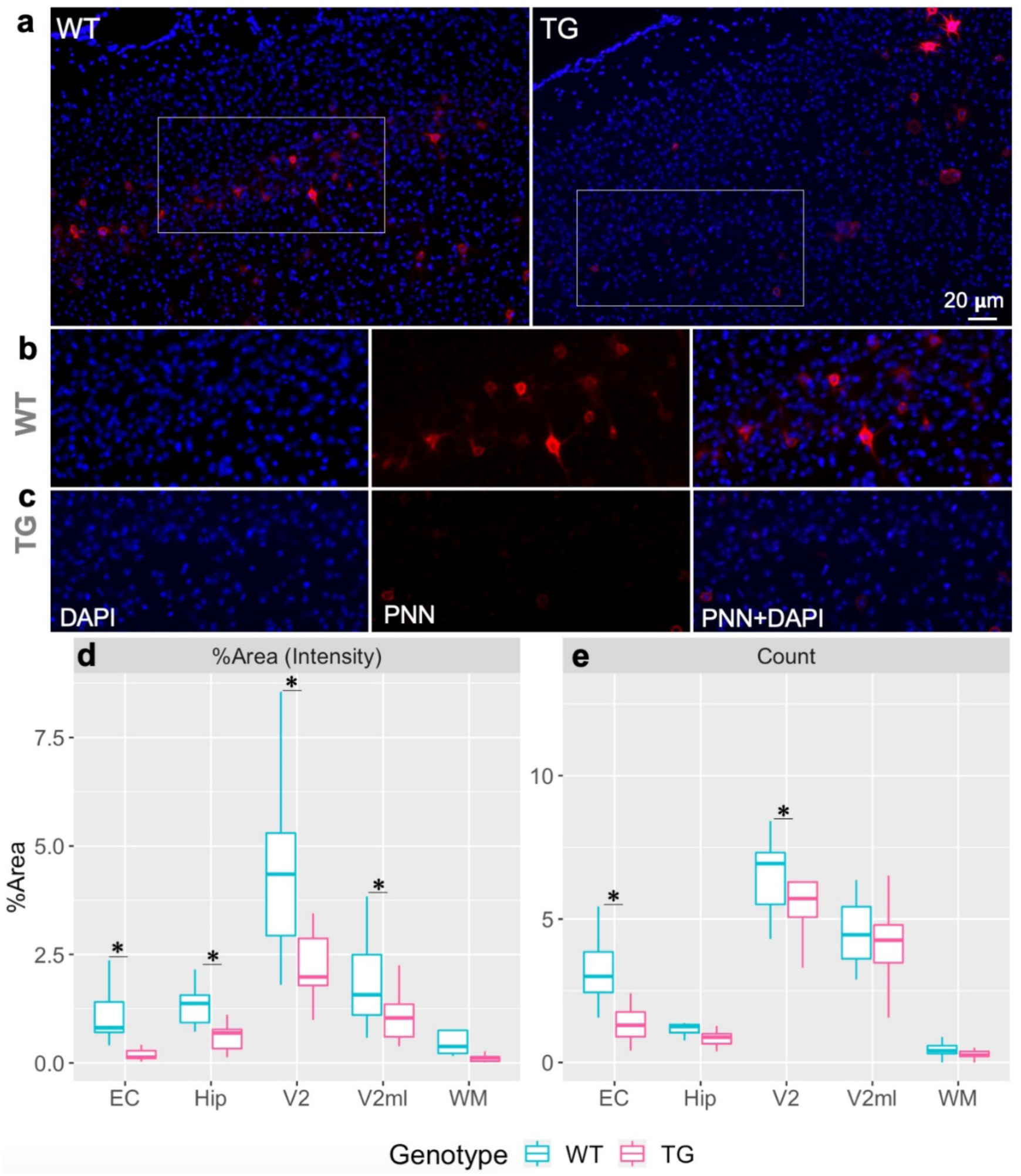
Transgenic mice had a reduced PNN intensity in the brain regions that show disrupted connectivity (d). In addition, PNN count reduced in the entorhinal cortex (EC), and secondary visual cortex l (V2) (e). Perineuronal nets (Red), DAPI-labelled nuclei (Blue), Double immunostained PNN and DAPI (Red + Blue). Mean % area intensity of PNN immunostaining (**d**) and mean PNN count (**e**), values are means ± SEM, *n* = 6/group. \**p* < 0.05, \*\**p* < 0.01, \*\*\**p* < 0.001 compared with the wild type (WT).

## Discussion

Our results of whole-brain fixel-based analysis demonstrated a reduced fiber density and fiber cross-section in the ArcAβ transgenic mice. We applied two complementary computational analyses: whole-brain structural connectivity and graph theory. Whole-brain structural connectivity analysis revealed a network disruption centered in the EC; while graph theoretical analysis identified decreased network properties. We also found a higher neurofilament light levels and a lower perineuronal net fluorescent intensity in the EC.

Using fixel-based analysis, we measured fiber tract-specific properties in the whole brain. FBA is a useful technique to measure microstructural alterations, such as fiber density and macrostructural changes including fiber bundle cross section. To our knowledge, this is the first demonstration of fiber tract-specific alterations in mouse model of Aβ amyloidosis. We showed reduced fiber density, fiber bundle cross section and fiber density plus bundle cross section in several brain regions (Fig. 2). The reduction of fiber tract-specific properties was shown in the right hemisphere, specifically in the corpus callosum, hippocampal commissure, retrosplenial cortex, dorsal hippocampus, and thalamus. Fiber tract-specific properties changes are also seen in the parahippocampal, cingulum bundle and fronto-occipital fasciculus in AD (Mito et al., 2018). Decreased fiber properties indicates a reduction in axons within a particular fiber bundle or a decreased intra-axonal volume of fibers. An alternative explanation is a reduction of the cross-sectional area of a fiber tract, resulting in smaller axonal diameter. Since we show that fiber density plus bundle cross section is reduced in TG mice, both are possible. However, it remains to be seen whether these weakened fiber tracts in TG mice would alter the global structural connectivity.

Whole-brain connectivity analysis revealed 5 nodes that displayed a disruption in structural connectivity: EC, subiculum, periaqueductal grey, lateral and mediolateral secondary visual cortices. The EC and hippocampus are two key vulnerable brain regions in AD, becoming affected in the early stages of AD (Mak et al., 2017, Hett et al., 2019). Disrupted structural connectivity in the EC is also a feature of AD (Mallio et al., 2015). Consistent with the human findings, we also showed structural connectivity deficits in the EC and hippocampal subfields, namely the subiculum. The subiculum is a part of the hippocampus, which carries outputs from the hippocampus to higher brain regions. Notably, entorhinal connectivity with higher cortical regions and medial temporal lobe structure contributes to cognition. EC provide output to the subiculum (O’Mara, 2005), while receives inputs from the sensory cortex including the visual cortex (Kerr et al., 2007). Overall, disrupted entorhinal connectivity could be the basis of cognitive deficit previously reported in TG mice (Knobloch et al., 2007b). Interestingly, the EC was identified as the largest module (based on the highest number of edges (*n* = 3)) of the disrupted network in TG mice. It is possible that EC neurons are more vulnerable to the amyloid deposition compared to the other brain regions (Harris et al., 2010). Collectively, our results of structural connectivity deficit might be the basis of decreased functional connectivity in the TG mice (Grandjean et al., 2016).

The connectome is a unique property, consistent across multiple species (van den Heuvel et al., 2016) and thus has important analytical value in translational research. Connectomics provide valuable information regarding the efficiency of information processing and topological organization (Dai and He, 2014). Here, we observed reduced local and global efficiency and degree centrality. A decreased local (Zhou and Lui, 2013) and global (Martensson et al., 2018) efficiency is a common characteristic of patients with AD. In addition, Aβ mouse model of amyloidosis (5XFAD mice) also showed a decreased network efficiency (Kesler et al., 2018). It is possible that axonal injury alters network properties. In fact, reduced myelination (Desai et al., 2010, Desai et al., 2009) and higher NFL release (Menke et al., 2015) may leave neurons susceptible to axonal injury. It is also possible that degradation of ECM causes deficits in neuronal connectivity (Bikbaev et al., 2015).

A possible explanation of reduced fiber density and fiber bundle cross section may be due to increased levels of NFL we showed in this study. Existing evidence from human studies strongly support the relationship between raised NFL levels and WM microstructure damage (Moore et al., 2018, Osborn et al., 2018). Furthermore, several studies describe association between cognitive impairment and increased CSF (Zetterberg et al., 2016) and plasma NFL (Lin et al., 2018) in AD. Study on three different mouse models of neurodegenerative diseases (A53T-αS, P301S-Tau and APP/PS1) showed increased plasma and CSF NFL with age (Bacioglu et al., 2016). Surprisingly, NFL gene deletion increased amyloid plaque load and synaptic pathology in APP/PS1 TG mice (Fernandez-Martos et al., 2015). However, none of these studies quantified NFL in axons and dendrites. In summary, increased NFL levels may be the underlying reason of fiber properties in ArcAβ TG mice. We sought to answer an important question: how does NFL expression change with age and amyloidosis? The Aβ load at 6 months is low in TG mice compared to 11 months (Grandjean et al., 2014). CSF NFL increased either in parallel or prior to Aβ accumulation in the mouse model of AD (Zetterberg, 2016). Therefore, we measured NFL intensity at 6 months of age. Bacioglu and colleauges showed increased CSF NFL in P301S-tau, APPPS1 and A53T-αS prior to symptomatic onset (Bacioglu et al., 2016). The authors showed 10-1000-fold higher CSF NFL in these TG mice, whereas, we found approximately 100-fold increase in NFL in ArcA*β* TG mice. Taken together, reduced fiber properties may due to the increased NFL in TG mice.

To further investigate the underlying basis of connectivity disruption, we measured the expression of MBP. MBP is the major constituent of brain myelin and is synthesized by oligodendrocyte and Schwann. A*β* peptide is cytotoxic to oligodendrocyte cells, acting through sphingomyelinase-ceramide signaling (Lee et al., 2004). However, our study reveals no demyelination in the brain regions that show disrupted connectivity. The possible explanations for this result are the selection of myelin marker. We measured the expression of MBP, but did not measure the expression of other myelin proteins, such as myelin proteolipid protein (MPP) and myelin oligodendrocyte glycoprotein (MOG). It is unknown whether the APP arctic mutation has an impact on MPP or MOG.

Connectivity depends on many factors, such as synaptic density and neuronal wiring. Although we did not measure the pre- or post-synaptic density in the brain, a severe synaptic plasticity impairment in ArcAβ TG mice has been documented (Knobloch et al., 2007a). Importantly, PNN regulate the synaptic plasticity (Kochlamazashvili et al., 2010) and rewiring the network (Pollock et al., 2014) of the interneurons (Wen et al., 2018). In this regard, we hypothesized that disrupted connectivity might be due to the reduced PNN levels. We showed a reduced PNN intensity in all of five brain regions, while PNN count reduced in the EC and V2 region but not in the hippocampus in six months old TG mice.

PNNs wrap around the interneurons in the brain. The interneurons have complex dendritic arborization to regulate the activity of principal neurons. PNN has an indirect role over principal neurons via regulating synaptic plasticity (Sigal et al., 2019) and dendritic arborization (Stolp et al., 2019) of the interneuron. In addition, PNN plays a key role in retaining neuronal connectivity (Bikbaev et al., 2015). There is some inconsistency on PNN findings in the transgenic mouse model of AD. For example, some studies argued that the degradation of PNN restores memory in P301s tau mice (Yang et al., 2015) and reduces plaque burden in APPswe/PS1dE9 mice while others demonstrated unaltered PNN in Tg2576 mice (Morawski et al., 2010b). On the contrary, a number of studies suggest protective role of PNN. For instance, neurons having intact PNN never deposits neurofibrillary tangles (Morawski et al., 2012); PNN prevents Aβ-induced neurotoxicity in cortical neurons (Miyata et al., 2007) and neurons with aggrecan-linked PNN protect tau pathology (Morawski et al., 2010a). Furthermore, reduced PNN expression had recently shown in Tg2576 mice (Cattaud et al., 2018). In summary, most of the previous studies are in support of beneficial role of PNN in AD mouse model. Here, we first time provide compelling evidence of reduced PNN in ArcAβ TG mice. We suggest that inadequate PNN may be the possible explanation for disrupted structural connectivity.

Although the EC has been widely studied for its critical role in AD, its role in connectome deficits in mice has received less attention. We reveal EC-centered connectivity deficits in the ArcAβ TG mice. The vulnerability of the entorhinal neurons has been reported in many previous studies (Adams et al., 2019, Rodriguez et al., 2019, Braak and Del Tredici, 2020). Differential gene expression in the EC is a possible explanation for PNN degradation (Santa-Maria et al., 2010). Nevertheless, further studies are required to explore the genetic basis of PNN reduction in ArcAβ TG mice.

This study has some limitations, mainly that we were unable to include the MRI brain samples in our histology experiment, as those brain samples were exhausted in a previous study (Grandjean et al., 2016). In addition, the diffusion-weighted images had a voxel size of 0.15 × 0.13 × 0.45 mm. Therefore, one can argue that the crossing-fiber calculation was not realistic specially with 0.45 mm slice. However, this limitation was resolved by the application of probabilistic tractography and an optimum diffusion direction (36) (Fig 1). We did not measure MPP or MOG expression. Future study can quantify the expression of MPP and MOG. However, we, for the first time provided pieces of evidence of fiber tract-specific properties reduction in an Aβ mouse model of amyloidosis. In addition, we provide compelling evidence that EC-centered connectivity disruption associated with increased NFL and decreased PNN.

## Conclusion

Our study indicates an EC-specific vulnerability in ArcAβ TG mice. Whole-brain connectivity analysis revealed network disruption centered at the EC consistent with PNN and NFL. Higher levels of NFL expression demonstrate a possible link with reduced fiber density and fiber bundle cross section. In addition, disruption of structural connectivity might be a consequence of both increased NFL and reduced PNN intesnity in the EC. The EC and hippocampus are the primary brain regions that are affected in the early stages of AD. Although we found a structural alteration in the EC, there were no changes in the hippocampus. The discrepancies in these findings could be partly because of the heterogeneous vulnerability of PNN to amyloid toxicity in distinct brain regions, reinforcing the importance of considering specific brain regions in the study of Alzheimer’s pathology.

## Acknowledgement

This work was, in part, supported by grants from NIH (R01AG054102, R01AG053500, R01AG053242, and R21AG050804). In addition, we acknowledge the funding from Indiana University (Strategic Research Initiative fund, Precision Health Initiative fund, and P. Michael Conneally Professorship). We also thank the contribution of Dr. Stephanie Philtjens, Dr. Luke Child Dabin, Dom James Acri and Mason Douglas Tate for providing us valuable feedbacks while preparing this manuscript.

